# Amplification and extraction free quantitative detection of viral nucleic acids and single-base mismatches using magnetic signal amplification circuit

**DOI:** 10.1101/2022.12.24.521858

**Authors:** Enja Laureen Rösch, Rebecca Sack, Mohammad Suman Chowdhury, Florian Wolgast, Meinhard Schilling, Thilo Viereck, Aidin Lak

## Abstract

Established nucleic acid detection assays require extraction and purification before sequence amplification and/or enzymatic reactions, hampering their widespread applications in point-of-care (POC) formats. Magnetic immunoassays based on magnetic particle spectroscopy and magnetic nanoparticles (MNPs) are isothermal, extraction- and purification-free, and can be quantitative and benchtop, making them suitable for POC settings. Here, we demonstrate a Magnetic signal Amplification Circuit (MAC) that combines specificity of toehold-mediated DNA strand displacement with magnetic response of MNPs to a clustering/declustering process. Our MAC assays require neither amplification nor extraction of target nucleic acids, and reveal four times better sensitivity than that of a magnetic circuit without signal amplification. Using MAC, we detect a highly specific 43 nucleotides sequence of SARS-CoV-2 virus. The MAC enables sensing both DNA and RNA targets with varying lengths and resolving single-base mismatches. Our MAC can be a powerful tool for translating research of nucleic acids detection to the clinic.

## INTRODUCTION

Highly specific and sensitive detection of nucleic acids-based disease biomarkers such as pathogen DNA/RNA, circulating tumor DNA, and microRNA has gained enormous attention in the past few decades.^1–4^ While reverse-transcription polymerase chain reaction (RT-PCR) remains the gold standard for viral DNA/RNA detection, PCR requires extensive and expensive sample processing, well-equipped laboratories, and trained personal, and thus is not compatible with point-of-care (POC) settings. Classical bulk PCR relies on sequence amplification and is often less sensitive when it comes to detecting rare mutations and alleles mostly because of PCR inhibitors and less efficient amplification.^5^ Compartmentalization of PCR sample into picolitre volume droplets, using cutting-edge droplet microfluidics, significantly isolates PCR inhibitors from the target sequence before the thermal cycling starts and facilitates the detection of rare mutations and alleles.^6^ Nevertheless, droplet microfluidics demand expensive instrumentations and consumables including microfluidic chips, droplet stabilizers, and droplet destabilizers.^7^ Alternative amplification-based detection strategies including loop-mediated isothermal amplification (LAMP),^8–11^ rolling circle amplification (RCA),^12–14^ and a combination of both ^15,16^ have emerged over the years, enabling detection of a few hundred copies of viral RNA. Despite being sensitive, LAMP-based assays require multistep sample handling and high-temperature incubation 55-65°C, limiting their POC implementation.

Plasmonic and optical detection concepts have significantly pushed the sensitivity of nucleic acid assays, yet these assays are not easily adaptable to detect the target in turbid biological samples due to signal attenuation.^17^ Moreover, the complexity of optical systems makes their miniaturization towards POC formats very challenging. Nanoparticle-based sensing platforms have recently taken major steps towards high sensitivity, yet by employing complex read-out system.^18^ They still fall short if detection needs to be highly specific and quantitative. Magnetic nanoparticles (MNPs) offer a highly promising sensing platform by harnessing their magnetic relaxation dynamics being highly sensitive to molecular interactions between the receptors on MNPs and targeting analytes.^19–22^ Magnetic immunoassays (MIAs) are rapid, isothermal, extraction- and purification-free, inherently quantitative, and can directly be performed on turbid bodily fluids due to no signal attenuation.^23^ Cryogenic superconducting quantum interference device (SQUID) magnetometers^20^ and nuclear magnetic resonance (NMR/MRI) readers^24,25^ have been used in early studies to detect nucleic acids. However, these are quite sophisticated and expensive approaches in terms of instrumentation.

The group of Weissleder has proposed “magnetic relaxation switches” as diagnostic magnetic resonance (DMR) for detecting proteins, enzymatic activity, viruses, and nucleic acids.^21,24^ Since then, alternative measurement modes including direct current (DC) magnetorelaxometry,^26–29^ alternating current (AC) susceptometry,^30–33^ and rotating magnetic fields^34,35^ have been successfully used to advance MIAs. Significant improvements regarding the assay sensitivity and miniaturization came when magnetic particle spectroscopy (MPS) has been transformed into a highly sensitive and cost-effective technique.^36–40^ Using the MPS-MIA sensing platform, detection of mimic virus particles, viral proteins, and pathogen specific nucleic acids have been demonstrated, where the assays rely on clustering of MNPs upon sensing the target.^41–43^ However, the clustering assays suffer from unspecificity, as MNPs are highly prone to unspecific clustering through magnetic, electrostatic, and van der Waals interactions.^44^ Consequently, the assay specificity is largely compromised. Remarkably and despite exclusive features, MIAs have not yet been successful in POC settings. Indeed, different assay designs are needed to fully exploit the benefit of MIAs. DNA has been widely used to build clusters of different nanoparticles due to its high programmability and specificity.^45–47^ In addition, the DNA switches had been revolutionized when the concept of toehold-mediated (TM) strand displacement (DSD) was first introduced by Yurke et al.^48^ A reversal DNA hybridization process based on several DNA walkers and robots have also been investigated. Being highly specific, molecular biosensors employing TM-DSD switches have gained numerous attentions in the past few years for enzyme-free amplification and detection of nucleic acids^49–52^ as well as detecting mismatches in targeting sequences.^53^

Here, we demonstrate a MIA platform that combines the features of TM-DSD and declustering of MNP-clusters. This platform requires no amplification and extraction of target nucleic acids, yet it achieves high sensitivity and specificity. Our so-called magnetic signal amplification circuit (MAC) enables amplification of magnetic signals, occurring upon declustering of MNP-clusters into MNPs. The declustering process is initially triggered by the binding of the target nucleic acids to the toehold regions and further accelerated by the release of amplification strands. Consequently, the MAC goes through several cycles of magnetic signal amplification, here called domino effect. Thus, MNPs act as proxies to detect the target nucleic acids. We exploit the MAC to detect highly specific SARS-CoV-2 nucleic acid sequences. We reveal that the MAC has four times better limit-of-detection (LOD) than that of a magnetic circuit (MC) without signal amplification. Significantly, the MAC resolves a single nucleotide mismatch in a 43 nt long DNA sequence. Additionally, it enables detecting different lengths of both DNA and RNA sequences in low picomolar range directly in virus lysis buffer. Our MAC detection platform opens the possibility towards combining the high specificity of nucleic acid targets with the enzyme-, amplification-, and extraction-free features of magnetic biosensing.

## RESULSTS AND DISCUSSION

Current clustering-based MIAs have reached the LOD in low nM range for detection of DNA^54^ and viral proteins,^55^ yet insufficient for testing clinical samples. Unlike other immunoassays,^56^ to reach the desired sensitivity, adaptability, and specificity that can push the MIAs towards clinical applications, we developed a facile declustering-based MIAs; where magnetic clusters dissociate into MNPs in presence of target nucleic acids. Therefore, unspecific clustering of nanoparticles does not interfere with the assay result (Fig. 1a). To improve the sensitivity, we designed magnetic signal amplification circuit (MAC) by integrating TM-DSD molecular switches into responsive magnetic clusters (RMCs) (Fig. 1b). The magnetic signal amplification is achieved through magnetic relaxation of MNPs, where the characteristic frequency of the Brownian magnetization relaxation process and concomitantly the MPS harmonics amplitude increase,^39^ while RMCs start declustering into MNPs upon hosting the input target sequence (Fig. 1a, schematic MPS spectra).

**Figure 1.**
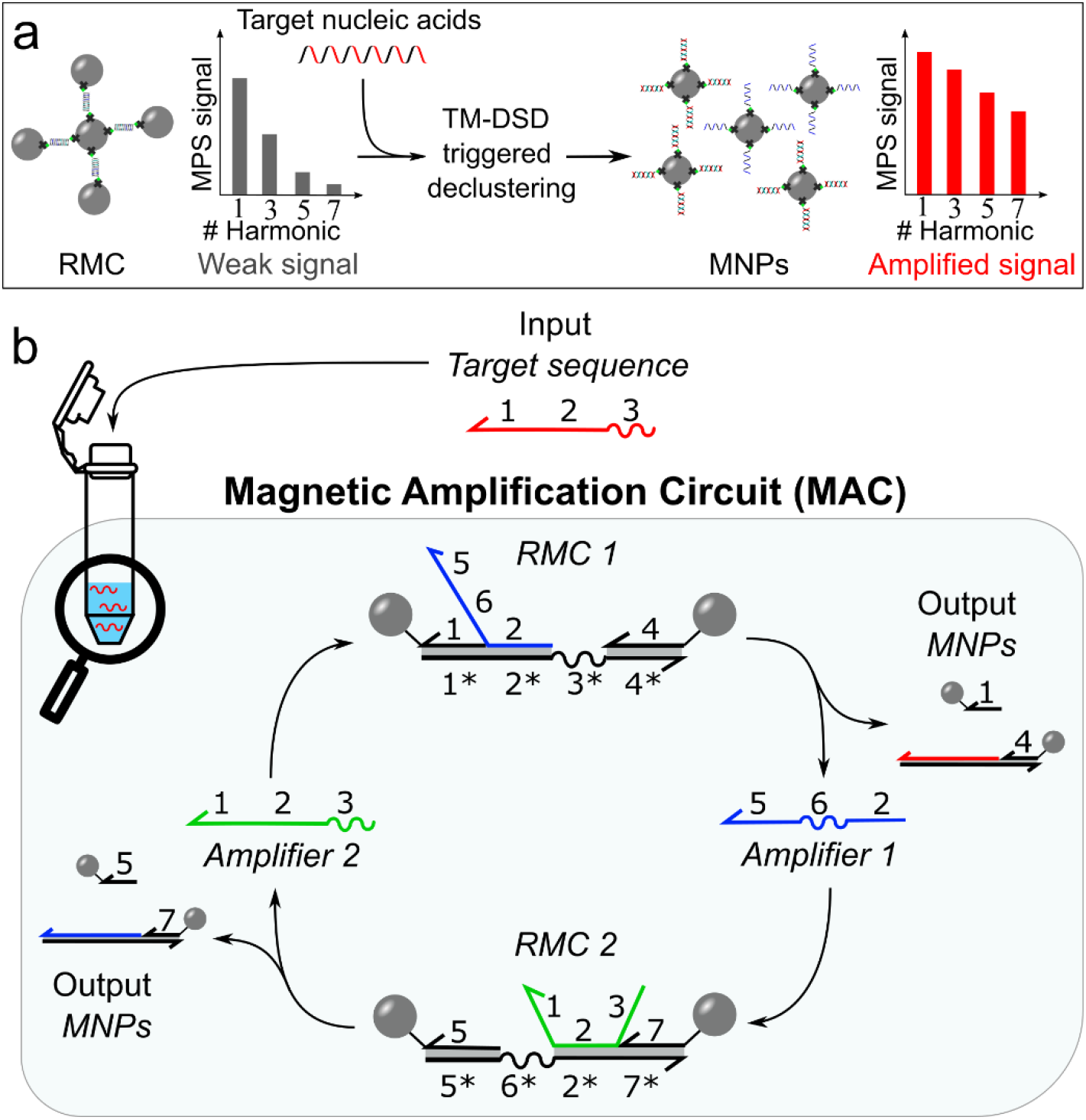
Working principle of declustering-based MAC nucleic acid sensing platform. (a) Schematic illustration of MPS magnetic signal amplification through the declustering of magnetic clusters into MNPs. DNA-labelled MNPs are tethered into responsive magnetic clusters (RMC) by the probe DNA construct. First, the input target sequence binds to the toehold region of the probe DNA, triggering the DSD reaction. The RMCs are subsequently declustered into output MNPs, which leads to an amplified MPS signal. (b) The MAC circuit consists of two different RMC systems: RMC1 and RMC2. A RMC system is formed by mixing the corresponding probe DNA already annealed with the amplifier sequences at the domain 2* with ssDNA labelled BNF-MNPs. The input nucleic acid target binds first to the toehold region 3* of RMC1. The displacement of the domain 2 and 1 by the target through a DSD reaction releases the amplifier 1 and generates the output MNPs. The amplifier 1, when released, binds to the toehold 6* of RMC2 and triggers the declustering of RMC2 by the DSD displacement of domain 5 and release of amplifier 2. Next, the amplifier 2, that is the DNA analog of the target nucleic acid, binds again to the toehold region of RMC1 and closes the cycle.

The MAC assay test tube consists of two RMC systems, namely RMC1 and RMC2 (see SI for the experimental procedures). A typical MAC assay is initiated when the target sequence is added into a one-to-one mixture of RMC1 and RMC2 (Fig. 1b). The probe DNA on the RMC1 possesses a 7 nt long toehold (3*), that is complementary to the section 3 of the target sequence. Next, the target nucleic acid binds to the toehold region and initiates the first DSD reaction through displacing the domains 2 and 1, thus releasing the amplifier sequence 1 and leading to declustering of the RMC1 into single MNPs. The amplifier sequence 1 possesses a domain complementary to the 7 nt toehold region of the probe DNA on the RMC2 (6*). Upon a binding event, the RMC2 clusters start declustering into single MNPs and thus releasing the amplifier sequence 2. The amplifier 2 is in fact the DNA analog of the target nucleic acid, and enables the next DSD reaction on the RMC1 to occur and thus closes one signal amplification cycle. Our MAC circuit exploits a so-called domino effect, that is realized by passing the amplifier sequences between the two RMC systems, which pushes the MAC circuit into a continuous cycle of MNP release and thus the assay read-out signal gets amplified.

In this study, BNF80 (micromod Partikeltechnologie GmbH, Rostock, Germany) MNPs were used as building blocks of RMCs (see Fig. S1 for full characterization of BNF-MNPs). Using dynamic light scattering (DLS), we performed particle size distribution (PSD) analysis on RMCs at pre- and post-MAC stages. We observed that the PSD at the post-MAC stage goes back to the PSD of the building blocks used for RMCs, which indicates a successful target and MAC-triggered declustering of RMCs (Fig. 2a). Additionally, to see the change in harmonics spectra at the pre- and post-MAC stages, we conducted MPS analyses that showed a weak harmonics spectrum at the pre-MAC stage, suggesting a slow Brownian relaxation of the RMCs. Upon adding the target sequence as an input, the post-MAC MPS spectrum resembled the one recorded for the building blocks of RMCs (Fig. 2b), demonstrating an efficient MAC-triggered declustering of RMCs. Furthermore, while looking at the MNPs, pre-, and post-MAC samples under scanning transmission electron microscope (STEM), we observed the formation of micrometer sized RMCs at pre-MAC stage that effectively declustered into nanometer sized MNPs upon binding to the target and going through the MAC circuit (Fig. 2c-e).

**Figure 2.**
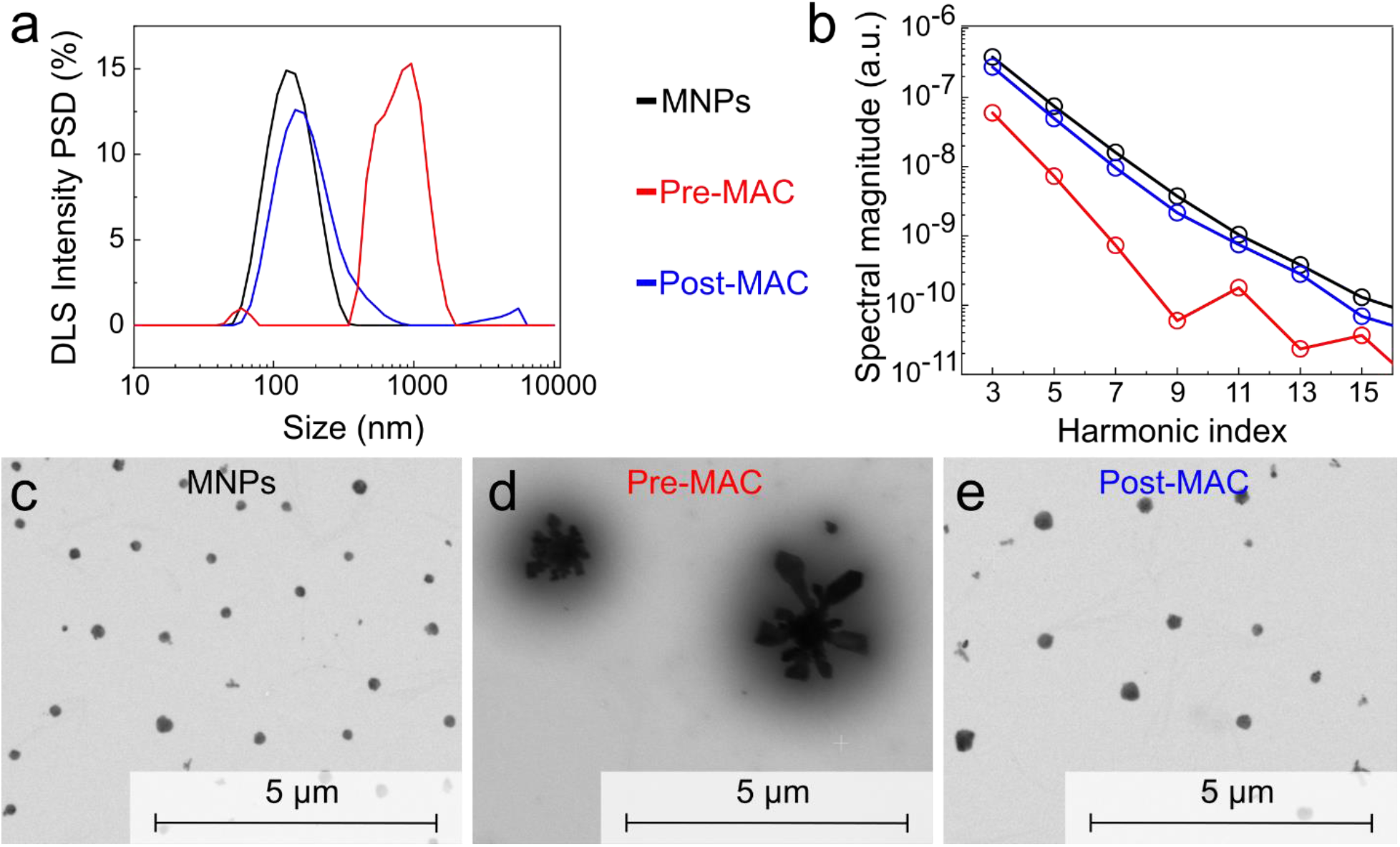
Colloidal, magnetization dynamics, and morphological changes upon forming RMCs (pre-MAC) and MAC-triggered declustering (post-MAC). Dynamic light scattering (DLS), magnetic particle spectroscopy (MPS), and scanning transmission electron microscope (STEM) analyses of MNPs, pre-MAC (referring to RMCs), and post-MAC (referring to declustered RMCs) samples. (a) The intensity PSD results measured by DLS show how the hydrodynamic size of post-MAC clusters switches back to the PSD of the building blocks used for RMCs. (b) The MPS spectra of single MNPs, pre-, and post-MAC samples reveal the intensification of the MPS signal upon sensing the target in post-MAC sample. Same color coding is applied to the panels (a) and (b). (c-e) STEM micrographs of single MNPs, pre-, and post-MAC samples showing how RMCs decluster back to single building blocks MNPs after MAC.

This motivated us to first investigate to what extent the assay LOD benefits from the MAC principle. We, therefore, focused on highly specific DNA analog of ORF1b gene of the SARS-CoV-2 genome (DNA/RNA sequences are given in table S1 and S2). The target sequence, derived from primers designed by Lopez-Rincon et al.,^57^ was adapted to our construct and extended to 43 nt (Fig. 3a). We performed the MAC assays on this specific 43 nt DNA sequence at different target concentrations and measured their MPS harmonics spectrum with our custom-built immunoMPS spectrometer (see assay protocols in SI).^39^ To eliminate the effect of particle concentration on the assay result, we used MPS H5^th^/H3^rd^ (HR53) harmonics ratio as the read-out indicator.^58^ Plotting the MPS HR53 ratio versus target concentration, a so-called dose-response plot is created, which shows that the HR53 increases steeply with the target DNA concentrations and saturates at 1.5 nM DNA concentration (Fig. 3b, normalized curves are shown in Fig. S2). This can be better seen by looking at the dose-response curves shown with 0.5 nM DNA concentration (Fig. 2c). The LOD of 27 pM (corresponding to 2 fmol) was determined by applying the 3sd criterion of control sample having two RMC systems without any target. Witnessing an order of magnitude better LOD of the MAC assays compared to the clustering MIAs i.e. ≈ 220 pM,^54^ we decided to compare the MAC circuit with a non-amplifying magnetic circuit (MC) (see scheme of the MC circuit in Fig. S3). Looking at the dose-response curve of the non-amplifying MC, we observed a shallow and linear rise in the HR53 with the target concentrations (Fig. 3b, c). The MC-based assay reached LOD of 106 pM (corresponding to 8 fmol), justifying the necessity of MAC-based assay design for an improved assay sensitivity. To estimate the displacement efficiency, the HR53 ratio was normalized as follows: (*HR53*_c_-*HR53*_min_)/(*HR53*_max_-*HR53*_min_), where *HR53*_c_ is the harmonics ratio at a target concentration (*c*) and *HR53*_min_ and *HR53*_max_ are the harmonics ratios at 0 and 10 nM target concentrations, respectively. Notably, the displacement efficiency for the MAC assay at 1.5 nM is as high as 95.5 %, while only 49.8 % for the non-amplifying MC. We tested leakage of our MAC circuit by plotting the harmonics ratio of the control sample immediately after mixing two RMC systems at 0 nM target concentration (the lowest red circle in Fig. 3b). The leakage can be attributed to that fact that no washing step is applied after annealing the probe DNA construct, wherein the added ratio of amplifier to cDNA is 2:1, meaning a portion of unreacted amplifiers is leading the leakage.

**Figure 3.**
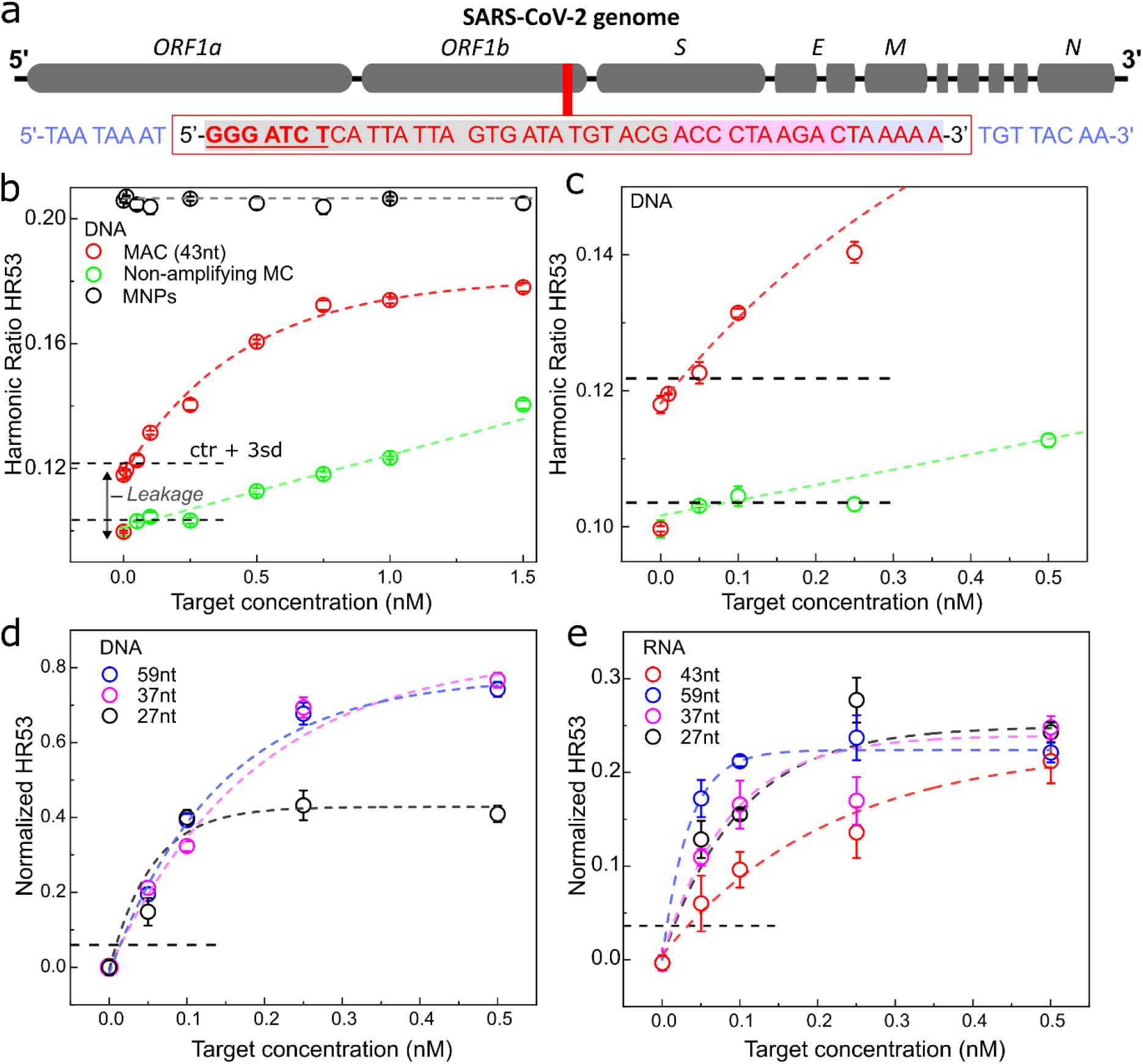
Detection of SARS-CoV-2 virus DNA analog and RNA of different lengths using MAC. (a) Region of the target sequence on the ORF1b gene from the SARS-CoV-2 viral genome. The 43 nt long sequence shown in the red box is the main target sequence. Additional nucleotides of the 59 nt sequence are colored in blue out of the red box. The shortened target sequences are shaded in light gray and light magenta color. (b) Harmonics HR53 ratio of both MAC (red) and non-amplifying MC (green) assays as a function of target concentration. The control sample is a mixture of both RMC systems in the absence of the target, and was measured right after the sample preparation and after 24 h incubation. The increase in the HR53 at 0 nM target concentration, here referred to as the leakage, is due to the presence of the amplifiers that are still present in the mixture as no cleaning step is done in between. As an additional control, we added the same target concentrations to a mixture of DNA-labelled BNF-MNPs, that were used as building blocks to form the RMCs, and observed no changes in the HR53 ratio for all target concentrations tested here. This result demonstrates that the magnetization dynamics of building block MNPs is not influenced by the target. (c) Shows the panel (b) zoomed in at the low concentrations. (d) MAC assays on DNA targets with different lengths. Normalized HR53 ratio vs. target concentration for MAC assays on 59 nt (blue), 37 nt (magenta) and 27 nt (black) DNA target sequences. (e) MAC assays on RNA target sequences with different lengths. Normalized HR53 harmonics ratio vs. target concentration for 59 nt (blue), 43 nt (red), 37 nt (magenta) and 27 nt (black) sequences. RNA assays were performed in the virus lysis buffer. All assays were incubated at 25°C for 24 h. Mean values and standard deviations (sd) were obtained from three independent measurements. Colored dashed lines are fits to the data points. The black horizontal dashed lines in all the panels mark the LOD cutoff line (control sample (mixture of two RMC systems at 0 nM target) + 3sd).

We next assessed the adaptability of our MAC assays. A true challenge for MAC assay is whether it enables sensing nucleic acids with not an optimal/designed length. Local conformations such as hairpins or pseudoknots strongly depend on the sequence length and bases,^59^ thus hindering the binding of target and limiting the assay sensitivity. Moreover, lysis of virus as well as other sample collection processes can break the RNA genome into fragments of random lengths, suggesting a specific length of the target sequence cannot always be guaranteed. Therefore, a robust assay should detect a target that is shorter and/or longer than the original probe DNA design.^60^ To address this, we performed assays on 27, 37, and 59 nt long target DNA sequences and compared with the original 43 nt long target sequence. The assays performed with these three sequences revealed dose-response patterns similar to what was observed for the 43 nt target, meaning a very steep initial rise of the HR53 ratio with the target concentration (Fig. 3d). The obtained LOD values of these three different sequences were also comparable to the LOD of 43 nt long sequence.

To qualify for the measurements of SARS-CoV-2 patient sample, the next logical step was to run MAC assays on RNA targets. For these studies, we took the same target sequences but as RNA. All RNA-based assays were performed on virus lysis buffer to mimic the environment of patient sampling (see SI for protocols). For RNA detection, our MAC offers a unique feature where it translates RNA assay to DNA binding reactions after the circuit is initially triggered by the target RNA. Thus, the assay becomes highly useful and robust for sensitive RNA samples. The dose-response data were recorded on all target lengths that were also tested in the DNA assays. The dose-response data shows a steep initial rise of the HR53 harmonics ratio with the target concentrations, which is similar to the read-outs of DNA-based assays (Fig. 3e). However, the LOD of RNA-based assay is 96 pM which is roughly three times the LOD of DNA-based assay, yet in the low pM regime, justifying that. The assay results on different RNA target lengths show that the length of the target does not compromise the performance of the MAC assay.

The TM-DSD reaction is highly sensitive to base-pair mismatches.^61–63^ We, therefore, decided to challenge our DNA-based MAC with base mismatches. To do so, we placed mismatches at different positions along the target sequence. We investigated DNA sequences with one (M1), two (M2), and six mismatches (M6). We plotted the HR53 harmonics ratio of the three mismatched targets and compared them with the DNA target having no mismatches (M0) (Fig. 4). Judging the assay result being positive or negative by comparing the HR53 ratio to the control sample+3sd, it can be appreciated that our MAC-based assays are capable of telling a target variant with a single-base mismatch apart from the fully matched one. Although the results shown here are from three independent assays, we have to however admit that our mismatch experiments is significant influenced by the position, base, and nearest neighbor of the nonmatching base that dictate the success of the single-base mismatch assays.

**Figure 4.**
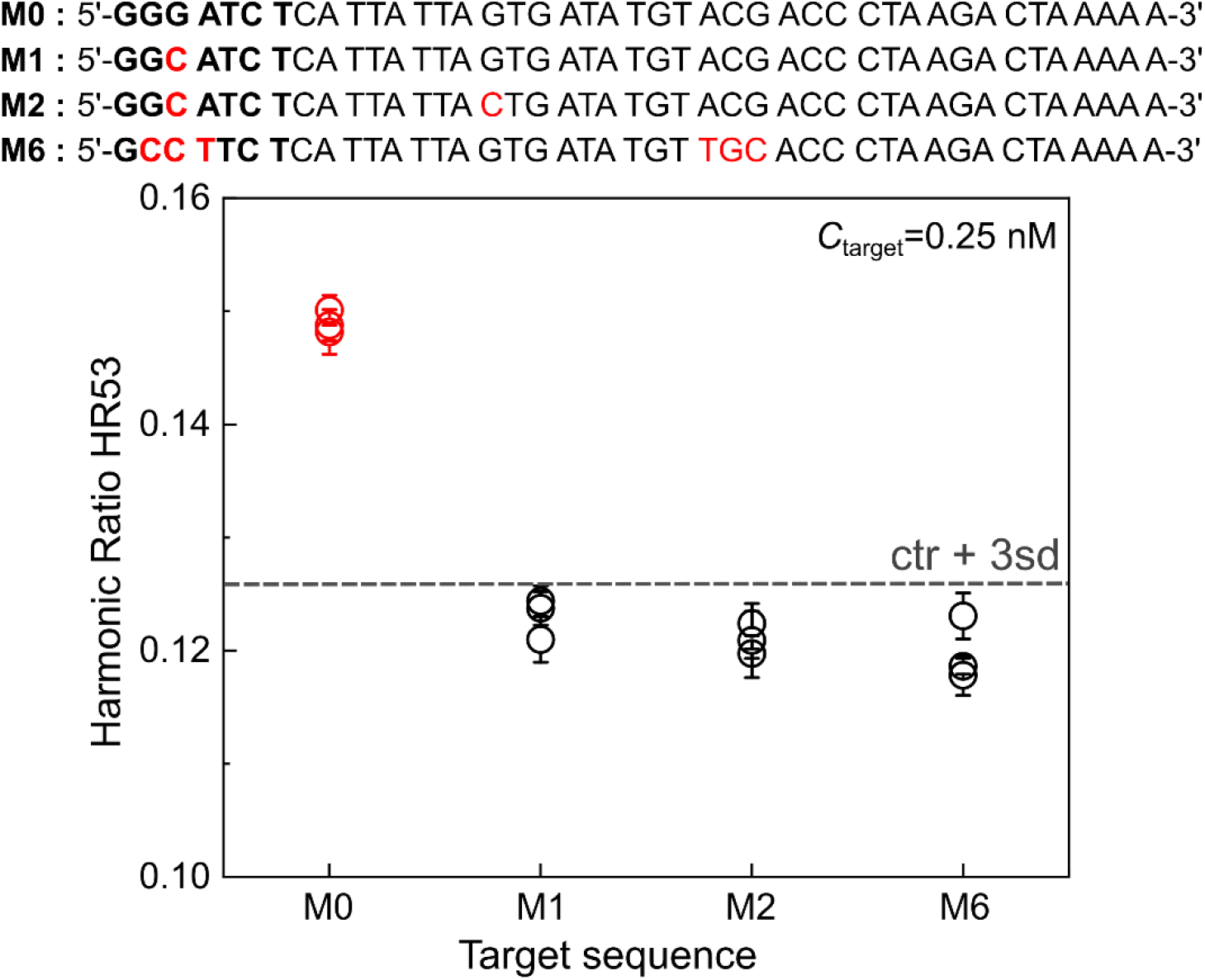
Specificity of MAC to nucleotide mismatches. Changes in the harmonics HR53 ratio for three mismatched DNA target sequences (M1-M2-M6) at a target concentration of 0.25 nM. For comparison, the change in the harmonics HR53 ratio for fully matching DNA target is plotted. The results of three independent assays are shown here. The dashed gray line marks the HR53 ratio the control sample (RMC1 + RMC2 + 0 nM target)+3d. The harmonics ratio was determined as described in the text.

### Conclusion

Here, we demonstrate magnetic signal amplification circuit (MAC) which works through declustering of responsive magnetic clusters and is responsive to the input of target nucleic acid. The MAC exploits the specificity of toehold-mediated DNA strand displacement and the magnetic response of MNPs to clustering/declustering. Additionally, the MAC benefits from the magnetic signal amplification through the domino effect based recycling of the amplifier sequences. Importantly, the MAC requires no amplification and extraction of target nucleic acids, yet it achieves high sensitivity and specificity.

Our MAC sensing concept addresses two major limitations of reported MIAs: sensitivity and specificity. In our assays, the MPS signal is amplified while sensing the target, which is in stark contrast to clustering-based MIAs, wherein the assay read-out is weakened upon detecting the target. We demonstrated that the MAC improves the LOD by four times, reaching to 27 pM (corresponding to 2 fmole), compared to the non-amplifying magnetic circuit. Moreover, we showed that the MAC circuit enables detection of different lengths of DNA and RNA targets directly in virus lysis buffer. Further studies on sequence mismatches revealed that the single-base mismatch resolution of the MAC assays can be achieved when the mismatch is positioned in the middle of the toehold region of the target.

The MAC circuit design can be adapted to DNA, RNA, and miRNAs. Its ability to resolve singe-base mismatch makes it highly suitable for mutant/variant testing. Working regardless of the target length, the MAC is highly compatible to testing patient samples, wherein a certain level of flexibility in terms of the length of DNA-RNA fragments being sought after is necessary. Being one-pot and purification-free, it facilitates translating magnetic biosensing into POC settings, where complex sample handling should be avoided. We expect to further improve the LOD by two to three orders of magnitudes by combining the domino effect with a circuit that offers an exponential amplification.

## Supporting information

Supporting Informtion

## ASSOCIATED CONTENT

Experimental details as mentioned in the test including: nucleic acid sequences, sample preparation protocols for DNA/RNA assays, and characterization techniques. Characterization of static and dynamic properties of BNF80 magnetic nanoparticles (Fig. S1), normalized data of harmonics ratio as a function of target concentration for DNA MAC and non-amplifying DNA MC assays (Fig. S2), scheme of non-amplifying MC circuit (Fig. S3), and MAC assay on 43nt RNA target up to 6 nM target concentration (Fig. S4).

## AUTHOR INFORMATION

### Present Addresses

### Author Contributions

A.L conceived the research idea. E.L.R. designed the research, prepared and characterized the MACs, performed the assays, analyzed the data, prepared the figures, and wrote the first draft of the manuscript. R.S. designed the research, characterized the MNPs, and prepared the assay samples. M.S.C wrote the manuscript. F.W. and T.V. designed and built the benchtop MPS spectrometer. M.S. provided research funding. A.L. designed the research, analyzed the data, supervised the study, and wrote the manuscript. All authors have given approval to the final version of the manuscript.

## ACKNOWLEDGMENT

This work is supported by DFG RTG 1952 “NanoMet”, Junior Research Group “Metrology4life”, and the add-on fellowship of Joachim Herz Foundation. We acknowledge Prof. Karsten Hiller (BRICS, TU Braunschweig) for providing biosafety Laboratory level 2 for RNA assays and Miss. Petra Schmidt (TU Braunschweig) for ICP-OES measurements.

## Notes

### Competing Interest Statement

The authors have declared no competing interest.

## REFERENCES

(1) Tavallaie, R.; McCarroll, J.; Le Grand, M.; Ariotti, N.; Schuhmann, W.; Bakker, E.; Tilley, R. D.; Hibbert, D. B.; Kavallaris, M.; Gooding, J. J. Nucleic Acid Hybridization on an Electrically Reconfigurable Network of Gold-Coated Magnetic Nanoparticles Enables MicroRNA Detection in Blood. Nat. Nanotechnol. 2018, 13 (11), 1066–1071. https://doi.org/10.1038/s41565-018-0232-x.

(2) Jiménez-Avalos, J. A.; Fernández-Macías, J. C.; González-Palomo, A. K. Circulating Exosomal MicroRNAs: New Non-Invasive Biomarkers of Non-Communicable Disease. Mol. Biol. Rep. 2021, 48 (1), 961–967. https://doi.org/10.1007/s11033-020-06050-w.

(3) Liu, T. Y.; Knott, G. J.; Smock, D. C. J.; Desmarais, J. J.; Son, S.; Bhuiya, A.; Jakhanwal, S.; Prywes, N.; Agrawal, S.; Díaz de León Derby, M.; Switz, N. A.; Armstrong, M.; Harris, A. R.; Charles, E. J.; Thornton, B. W.; Fozouni, P.; Shu, J.; Stephens, S. I.; Kumar, G. R.; Zhao, C.; Mok, A.; Iavarone, A. T.; Escajeda, A. M.; McIntosh, R.; Kim, S.; Dugan, E. J.; Hamilton, J. R.; Lin-Shiao, E.; Stahl, E. C.; Tsuchida, C. A.; Moehle, E. A.; Giannikopoulos, P.; McElroy, M.; McDevitt, S.; Zur, A.; Sylvain, I.; Ciling, A.; Zhu, M.; Williams, C.; Baldwin, A.; Pollard, K. S.; Tan, M. X.; Ott, M.; Fletcher, D. A.; Lareau, L. F.; Hsu, P. D.; Savage, D. F.; Doudna, J. A. Accelerated RNA Detection Using Tandem CRISPR Nucleases. Nat. Chem. Biol. 2021, 17 (9), 982–988. https://doi.org/10.1038/s41589-021-00842-2.

(4) Lu, W.; Chen, Y.; Liu, Z.; Tang, W.; Feng, Q.; Sun, J.; Jiang, X. Quantitative Detection of MicroRNA in One Step via Next Generation Magnetic Relaxation Switch Sensing. ACS Nano 2016, 10 (7), 6685–6692. https://doi.org/10.1021/acsnano.6b01903.

(5) Hindson, B. J.; Ness, K. D.; Masquelier, D. A.; Belgrader, P.; Heredia, N. J.; Makarewicz, A. J.; Bright, I. J.; Lucero, M. Y.; Hiddessen, A. L.; Legler, T. C.; Kitano, T. K.; Hodel, M. R.; Petersen, J. F.; Wyatt, P. W.; Steenblock, E. R.; Shah, P. H.; Bousse, L. J.; Troup, C. B.; Mellen, J. C.; Wittmann, D. K.; Erndt, N. G.; Cauley, T. H.; Koehler, R. T.; So, A. P.; Dube, S.; Rose, K. A.; Montesclaros, L.; Wang, S.; Stumbo, D. P.; Hodges, S. P.; Romine, S.; Milanovich, F. P.; White, H. E.; Regan, J. F.; Karlin-Neumann, G. A.; Hindson, C. M.; Saxonov, S.; Colston, B. W. High-Throughput Droplet Digital PCR System for Absolute Quantitation of DNA Copy Number. Anal. Chem. 2011, 83 (22), 8604–8610. https://doi.org/10.1021/ac202028g.

(6) Kojabad, A. A.; Farzanehpour, M.; Galeh, H. E. G.; Dorostkar, R.; Jafarpour, A.; Bolandian, M.; Nodooshan, M. M. Droplet Digital PCR of Viral DNA/RNA, Current Progress, Challenges, and Future Perspectives. J. Med. Virol. 2021, 93 (7), 4182–4197. https://doi.org/10.1002/jmv.26846.

(7) Chowdhury, M. S.; Zheng, W.; Kumari, S.; Heyman, J.; Zhang, X.; Dey, P.; Weitz, D. A.; Haag, R. Dendronized Fluorosurfactant for Highly Stable Water-in-Fluorinated Oil Emulsions with Minimal Inter-Droplet Transfer of Small Molecules. Nat. Commun. 2019, 10 (1). https://doi.org/10.1038/s41467-019-12462-5.

(8) Chaouch, M. Loop‐mediated Isothermal Amplification LAMP An Effective Molecular.Pdf. Rev Med Virol. 2021, 31, e2215.

(9) Dao Thi, V. L.; Herbst, K.; Boerner, K.; Meurer, M.; Kremer, L. P. M.; Kirrmaier, D.; Freistaedter, A.; Papagiannidis, D.; Galmozzi, C.; Stanifer, M. L.; Boulant, S.; Klein, S.; Chlanda, P.; Khalid, D.; Miranda, I. B.; Schnitzler, P.; Kräusslich, H. G.; Knop, M.; Anders, S. A Colorimetric RT-LAMP Assay and LAMP-Sequencing for Detecting SARS-CoV-2 RNA in Clinical Samples. Sci. Transl. Med. 2020, 12 (556). https://doi.org/10.1126/SCITRANSLMED.ABC7075.

(10) Rabe, B. A.; Cepko, C. SARS-CoV-2 Detection Using Isothermal Amplification and a Rapid, Inexpensive Protocol for Sample Inactivation and Purification. Proc. Natl. Acad. Sci. U. S. A. 2020, 117 (39), 24450–24458. https://doi.org/10.1073/pnas.2011221117.

(11) Ganguli, A.; Mostafa, A.; Berger, J.; Aydin, M. Y.; Sun, F.; Stewart de Ramirez, S. A.; Valera, E.; Cunningham, B. T.; King, W. P.; Bashir, R. Rapid Isothermal Amplification and Portable Detection System for SARS-CoV-2. Proc. Natl. Acad. Sci. U. S. A. 2020, 117 (37), 22727–22735. https://doi.org/10.1073/pnas.2014739117.

(12) Lizardi, P. M.; Huang, X.; Zhu, Z.; Bray-Ward, P.; Thomas, D. C.; Ward, D. C. Mutation Detection and Single-Molecule Counting Using Isothermal Rolling-Circle Amplification. Nat. Genet. 1998, 19 (3), 225–232. https://doi.org/10.1038/898.

(13) Choi, M. H.; Kumara, G. S. R.; Lee, J.; Seo, Y. J. Point-of-Care COVID-19 Testing: Colorimetric Diagnosis Using Rapid and Ultra-Sensitive Ramified Rolling Circle Amplification. Anal. Bioanal. Chem. 2022, 414 (19), 5907–5915. https://doi.org/10.1007/s00216-022-04156-7.

(14) Xu, L.; Duan, J.; Chen, J.; Ding, S.; Cheng, W. Recent Advances in Rolling Circle Amplification-Based Biosensing Strategies-A Review. Anal. Chim. Acta 2021, 1148. https://doi.org/10.1016/j.aca.2020.12.062.

(15) Tian, W.; Li, P.; He, W.; Liu, C.; Li, Z. Rolling Circle Extension-Actuated Loop-Mediated Isothermal Amplification (RCA-LAMP) for Ultrasensitive Detection of MicroRNAs. Biosens. Bioelectron. 2019, 128, 17–22. https://doi.org/10.1016/j.bios.2018.12.041.

(16) Hasegawa, T.; Hapsari, D.; Iwahashi, H. RNase H-Dependent Amplification Improves the Accuracy of Rolling Circle Amplification Combined with Loop-Mediated Isothermal Amplification (RCA-LAMP). PeerJ 2021, 9. https://doi.org/10.7717/peerj.11851.

(17) Qiu, G.; Gai, Z.; Tao, Y.; Schmitt, J.; Kullak-Ublick, G. A.; Wang, J. Dual-Functional Plasmonic Photothermal Biosensors for Highly Accurate Severe Acute Respiratory Syndrome Coronavirus 2 Detection. ACS Nano 2020, 14 (5), 5268–5277. https://doi.org/10.1021/acsnano.0c02439.

(18) Yu, Z.; Fang, W.; Yang, Y.; Yao, H.; Hu, P.; Shi, J. Non‐PCR Ultrasensitive Detection of Viral RNA by a Nanoprobe‐Coupling Strategy -1.Pdf. Adv Healthc. Mater. 2022, 11, 2200031.

(19) Kotitz, R.; Matz, H.; Trahms, L.; Koch, H.; Weitschies, W.; Rheinltinder, T.; Semmler, W.; Bunte, T. SQUID Based Remanence Measurements for Immunoassays. IEEE Trans. Appl. Supercond. 1997, 7 (2), 3678–3681.

(20) Grossman, H. L.; Myers, W. R.; Vreeland, V. J.; Bruehl, R.; Alper, M. D.; Bertozzi, C. R.; Clarke, J. Detection of Bacteria in Suspension by Using a Superconducting Quantum Interference Device. Proc. Natl. Acad. Sci. U. S. A. 2004, 101 (1), 129–134.

(21) Josephson, L.; Manuel Perez, J.; Weissleder, R. Magnetic Nanosensors for the Detection of Oligonucleotide Sequences. Angew. Chemie - Int. Ed. 2001, 40 (17), 3204–3206. https://doi.org/10.1002/1521-3773(20010903)40:17<3204::AID-ANIE3204>3.0.CO;2-H.

(22) Osterfeld, S. J.; Yu, H.; Gaster, R. S.; Caramuta, S.; Xu, L.; Han, S. J.; Hall, D. A.; Wilson, R. J.; Sun, S.; White, R. L.; Davis, R. W.; Pourmand, N.; Wang, S. X. Multiplex Protein Assays Based on Real-Time Magnetic Nanotag Sensing. Proc. Natl. Acad. Sci. U. S. A. 2008, 105 (52), 20637–20640. https://doi.org/10.1073/pnas.0810822105.

(23) Haun, J. B.; Yoon, T. J.; Lee, H.; Weissleder, R. Magnetic Nanoparticle Biosensors. Wiley Interdiscip. Rev. Nanomedicine Nanobiotechnology 2010, 2 (3), 291–304. https://doi.org/10.1002/wnan.84.

(24) Perez, J. M.; Josephson, L.; Weissleder, R. Use of Magnetic Nanoparticles as Nanosensors to Probe for Molecular Interactions. ChemBioChem 2004, 5 (3), 261–264. https://doi.org/10.1002/cbic.200300730.

(25) Lee, H.; Sun, E.; Ham, D.; Weissleder, R. Chip-NMR Biosensor for Detection and Molecular Analysis of Cells. Nat. Med. 2008, 14 (8), 869–874. https://doi.org/10.1038/nm.1711.

(26) Ludwig, F.; Heim, E.; Mäuselein, S.; Eberbeck, D.; Schilling, M. Magnetorelaxometry of Magnetic Nanoparticles with Fluxgate Magnetometers for the Analysis of Biological Targets. J. Magn. Magn. Mater. 2005, 293 (1), 690–695. https://doi.org/10.1016/j.jmmm.2005.02.045.

(27) Lange, J.; Kötitz, R.; Haller, A.; Trahms, L.; Semmler, W.; Weitschies, W. Magnetorelaxometry - A New Binding Specific Detection Method Based on Magnetic Nanoparticles. J. Magn. Magn. Mater. 2002, 252 (1-3 SPEC. ISS.), 381–383. https://doi.org/10.1016/S0304-8853(02)00657-1.

(28) Eberbeck, D.; Bergemann, C.; Wiekhorst, F.; Steinhoff, U.; Trahms, L. Quantification of Specific Bindings of Biomolecules by Magnetorelaxometry. J. Nanobiotechnology 2008, 6, 1–12. https://doi.org/10.1186/1477-3155-6-4.

(29) Enpuku, K.; Tokumitsu, H.; Sugimoto, Y.; Kuma, H.; Hamasaki, N.; Tsukamoto, A.; Mizoguchi, T.; Kandori, A.; Yoshinaga, K.; Kanzaki, H.; Usuki, N. Fast Detection of Biological Targets with Magnetic Marker and SQUID. IEEE Trans. Appl. Supercond. 2009, 19 (3), 844–847. https://doi.org/10.1109/TASC.2009.2018819.

(30) Chen, H.-H.; Hsu, M.-H.; Lee, K.-Hu.; Chen, W.-Y.; Yang, S.-Y. Real-Time Changes in the AC Magnetic Susceptibility of Reagents during Immunomagnetic Reduction Assays. AIP Adv. 2022, 12 (6), 065220. https://doi.org/10.1063/5.0094418.

(31) Tian, B.; Qiu, Z.; Ma, J.; Zardán Gómez de la Torre, T.; Johansson, C.; Svedlindh, P.; Strömberg, M. Attomolar Zika Virus Oligonucleotide Detection Based on Loop-Mediated Isothermal Amplification and AC Susceptometry. Biosens. Bioelectron. 2016, 86, 420–425. https://doi.org/10.1016/j.bios.2016.06.085.

(32) Fock, J.; Parmvi, M.; Strömberg, M.; Svedlindh, P.; Donolato, M.; Hansen, M. F. Comparison of Optomagnetic and AC Susceptibility Readouts in a Magnetic Nanoparticle Agglutination Assay for Detection of C-Reactive Protein. Biosens. Bioelectron. 2017, 88, 94–100. https://doi.org/10.1016/j.bios.2016.07.088.

(33) Mizoguchi, T.; Kandori, A.; Kawabata, R.; Ogata, K.; Hato, T.; Tsukamoto, A.; Adachi, S.; Tanabe, K.; Tanaka, S.; Tsukada, K.; Enpuku, K. Highly Sensitive Third-Harmonic Detection Method of Magnetic Nanoparticles Using an AC Susceptibility Measurement System for Liquid-Phase Assay. IEEE Trans. Appl. Supercond. 2016, 26 (5), 15–18. https://doi.org/10.1109/TASC.2016.2581703.

(34) Dieckhoff, J.; Lak, A.; Schilling, M.; Ludwig, F. Protein Detection with Magnetic Nanoparticles in a Rotating Magnetic Field. J. Appl. Phys. 2014, 115 (2). https://doi.org/10.1063/1.4861032.

(35) Lak, A.; Dieckhoff, J.; Ludwig, F.; Scholtyssek, J. M.; Goldmann, O.; Lünsdorf, H.; Eberbeck, D.; Kornowski, A.; Kraken, M.; Litterst, F. J.; Fiege, K.; Mischnick, P.; Schilling, M. Highly Stable Monodisperse PEGylated Iron Oxide Nanoparticle Aqueous Suspensions: A Nontoxic Tracer for Homogeneous Magnetic Bioassays. Nanoscale 2013, 5, 11447–11455.

(36) Gleich, B.; Weizenecker, J. Tomographic Imaging Using the Nonlinear Response of Magnetic Particles. Nature 2005, 435, 1214.

(37) Orlov, A. V.; Khodakova, J. A.; Nikitin, M. P.; Shepelyakovskaya, A. O.; Brovko, F. A.; Laman, A. G.; Grishin, E. V.; Nikitin, P. I. Magnetic Immunoassay for Detection of Staphylococcal Toxins in Complex Media. Anal. Chem. 2013, 85 (2), 1154–1163. https://doi.org/10.1021/ac303075b.

(38) Krause, H. J.; Wolters, N.; Zhang, Y.; Offenhäusser, A.; Miethe, P.; Meyer, M. H. F.; Hartmann, M.; Keusgen, M. Magnetic Particle Detection by Frequency Mixing for Immunoassay Applications. J. Magn. Magn. Mater. 2007, 311, 436–444. https://doi.org/10.1016/j.jmmm.2006.10.1164.

(39) Draack, S.; Lucht, N.; Remmer, H.; Martens, M.; Fischer, B.; Schilling, M.; Ludwig, F.; Viereck, T. Multiparametric Magnetic Particle Spectroscopy of CoFe2O4 Nanoparticles in Viscous Media. J. Phys. Chem. C 2019, 123 (11), 6787–6801.

(40) Vogel, P.; Rückert, M. A.; Friedrich, B.; Tietze, R.; Lyer, S.; Kampf, T.; Hennig, T.; Dölken, L.; Alexiou, C.; Behr, V. C. Critical Offset Magnetic PArticle SpectroScopy for Rapid and Highly Sensitive Medical Point-of-Care Diagnostics. Nat. Commun. 2022, 13 (1), 1–9. https://doi.org/10.1038/s41467-022-34941-y.

(41) Wu, K.; Su, D.; Saha, R.; Wong, D.; Wang, J. P. Magnetic Particle Spectroscopy-Based Bioassays: Methods, Applications, Advances, and Future Opportunities. J. Phys. D. Appl. Phys. 2019, 15 (17), 173001. https://doi.org/10.1088/1361-6463/ab03c0.

(42) Wu, K.; Liu, J.; Saha, R.; Su, D.; Krishna, V. D.; Cheeran, M. C. J.; Wang, J. P. Magnetic Particle Spectroscopy for Detection of Influenza A Virus Subtype H1N1. ACS Appl. Mater. Interfaces 2020, 12 (12), 13686–13697. https://doi.org/10.1021/acsami.0c00815.

(43) Zhong, J.; Rösch, E. L.; Viereck, T.; Schilling, M.; Ludwig, F. Toward Rapid and Sensitive Detection of SARS-CoV-2 with Functionalized Magnetic Nanoparticles. ACS Sensors 2021, 6 (3), 976–984. https://doi.org/10.1021/acssensors.0c02160.

(44) Durhuus, F. L.; Wandall, L. H.; Boisen, M. H.; Kure, M.; Beleggia, M.; Frandsen, C. Simulated Clustering Dynamics of Colloidal Magnetic Nanoparticles. Nanoscale 2021, 13 (3), 1970–1981. https://doi.org/10.1039/d0nr08561h.

(45) Mirkin, C. A.; Letsinger, R. L.; Mucic, R. C.; Storhoff, J. J. A DNA-Based Method for Rationally Assembling Nanoparticles into Macroscopic Materials*. Nature 1996, 382, 607–609. https://doi.org/10.4324/9780429200151-2.

(46) Nykypanchuk, D.; Maye, M. M.; Van Der Lelie, D.; Gang, O. DNA-Guided Crystallization of Colloidal Nanoparticles. Nature 2008, 451 (7178), 549–552. https://doi.org/10.1038/nature06560.

(47) Ryu, Y.; Jin, Z.; Lee, J.; Noh, S.; Shin, T.-H.; Jo, S.-M.; Choi, J.; Park, H.; Cheon, J.; Kim, H.-S. Size-Controlled Construction of Magnetic Nanoparticle Clusters Using DNA-Binding Zinc Finger Protein. Angew. Chemie Int. Ed. 2015, 54 (3), 923–926. https://doi.org/10.1002/anie.201408593.

(48) Yurke, B.; Turber, A. J.; Jr, A. P. M.; Simmel, F. C.; Neumann, J. L. ADNA-Fuelled Molecular Machine Made of DNA Bernard. Nature 2000, 406 (August), 605–608.

(49) Simmel, F. C.; Yurke, B.; Singh, H. R. Principles and Applications of Nucleic Acid Strand Displacement Reactions. Chem. Rev. 2019, 119 (10), 6326–6369. https://doi.org/10.1021/acs.chemrev.8b00580.

(50) Jung, J. K.; Archuleta, C. M.; Alam, K. K.; Lucks, J. B. Programming Cell-Free Biosensors with DNA Strand Displacement Circuits. Nat. Chem. Biol. 2022, 18 (4), 385–393. https://doi.org/10.1038/s41589-021-00962-9.

(51) Lv, H.; Li, Q.; Shi, J.; Fan, C.; Wang, F. Biocomputing Based on DNA Strand Displacement Reactions. ChemPhysChem 2021, 22 (12), 1151–1166. https://doi.org/10.1002/cphc.202100140.

(52) Oishi, M. Comparative Study of DNA Circuit System-Based Proportional and Exponential Amplification Strategies for Enzyme-Free and Rapid Detection of MiRNA at Room Temperature. ACS Omega 2018, 3 (3), 3321–3329. https://doi.org/10.1021/acsomega.7b01866.

(53) Wu, F.; Chen, M.; Lan, J.; Xia, Y.; Liu, M.; He, W.; Li, C.; Chen, X.; Chen, J. A Universal Locked Nucleic Acid-Integrated X-Shaped DNA Probe Design for Amplified Fluorescence Detection of Single-Nucleotide Variant. Sensors Actuators, B Chem. 2017, 241, 123–128. https://doi.org/10.1016/j.snb.2016.10.066.

(54) Rösch, E. L.; Zhong, J.; Lak, A.; Liu, Z.; Etzkorn, M.; Schilling, M.; Ludwig, F.; Viereck, T.; Lalkens, B. Point-of-Need Detection of Pathogen-Specific Nucleic Acid Targets Using Magnetic Particle Spectroscopy. Biosens. Bioelectron. 2021, 192, 113536. https://doi.org/10.1016/j.bios.2021.113536.

(55) Wu, K.; Chugh, V. K.; D. Krishna V.; di Girolamo, A.; Wang, Y. A.; Saha, R.; Liang, S.; Cheeran, M. C. J.; Wang, J. P. One-Step, Wash-Free, Nanoparticle Clustering-Based Magnetic Particle Spectroscopy Bioassay Method for Detection of SARS-CoV-2 Spike and Nucleocapsid Proteins in the Liquid Phase. ACS Appl. Mater. Interfaces 2021, 13 (37), 44136–44146. https://doi.org/10.1021/acsami.1c14657.

(56) Mata Calidonio, J.; Gomez-Marquez, J.; Hamad-Schifferli, K. Nanomaterial and Interface Advances in Immunoassay Biosensors. J. Phys. Chem. C 2022, 126 (42), 17804–17815. https://doi.org/10.1021/acs.jpcc.2c05008.

(57) Lopez-Rincon, A.; Tonda, A.; Mendoza-Maldonado, L.; Mulders, D. G. J. C.; Molenkamp, R.; Perez-Romero, C. A.; Claassen, E.; Garssen, J.; Kraneveld, A. D. Classification and Specific Primer Design for Accurate Detection of SARS-CoV-2 Using Deep Learning. Sci. Rep. 2021, 11 (1), 1–11. https://doi.org/10.1038/s41598-020-80363-5.

(58) Rauwerdink, A. M.; Weaver, J. B. Viscous Effects on Nanoparticle Magnetization Harmonics. J. Magn. Magn. Mater. 2010, 322 (6), 609–613. https://doi.org/10.1016/j.jmmm.2009.10.024.

(59) Dong, F.; Allawi, H. T.; Anderson, T.; Neri, B. P.; Lyamichev, V. I. Secondary Structure Prediction and Structure-Specific Sequence Analysis of Single-Stranded DNA. Nucleic Acids Res. 2001, 29 (15), 3248–3257. https://doi.org/10.1093/nar/29.15.3248.

(60) Chen, D.; Wu, Y.; Hoque, S.; Tilley, R. D.; Gooding, J. J. Rapid and Ultrasensitive Electrochemical Detection of Circulating Tumor DNA by Hybridization on the Network of Gold-Coated Magnetic Nanoparticles. Chem. Sci. 2021, 12 (14), 5196–5201. https://doi.org/10.1039/d1sc01044a.

(61) Machinek, R. R. F.; Ouldridge, T. E.; Haley, N. E. C.; Bath, J.; Turberfield, A. J. Programmable Energy Landscapes for Kinetic Control of DNA Strand Displacement. Nat. Commun. 2014, 5, 1–9. https://doi.org/10.1038/ncomms6324.

(62) Broadwater, D. W. B.; Kim, H. D. The Effect of Basepair Mismatch on DNA Strand Displacement. Biophys. J. 2016, 110 (7), 1476–1484. https://doi.org/10.1016/j.bpj.2016.02.027.

(63) Irmisch, P.; Ouldridge, T. E.; Seidel, R. Modeling DNA-Strand Displacement Reactions in the Presence of Base-Pair Mismatches. J. Am. Chem. Soc. 2020, 142 (26), 11451–11463. https://doi.org/10.1021/jacs.0c03105.

