## Supporting Informtion for "Amplification and extraction free quantitative detection of viral nucleic acids and single-base mismatches using magnetic signal amplification circuit"

**Materials and methods:**

**Chemicals:**

Bionized Nano Ferrites-Starch particles (BNF80) functionalized with streptavidin were purchased from micromod Partikeltechnologie GmbH (Rostock, Germany). The particle stock solution has an iron concentration of 5.5 $mg_{Fe}$/ml, corresponding to 20 nM particle concentration. TRIS-HCl and EDTA were purchased from Carl Roth. NaCl, Triton-X, Tween 20 and DTT were purchased from Sigma-Aldrich. RNase inhibitor was purchased from ThermoFisher Scientific. Carrier RNA was purchased from QIAGEN. No purification of the chemicals was carried out prior to their use. All DNA/RNA sequences were purchased from Eurofins Genomics (Germany). The sequences used are listed in Table S1. An Eppendorf ThermoMixer was used in all incubation/mix processes.

The Tris buffer is composed of 20 mM TRIS-HCl, pH 7.4, 1 mM EDTA, 150 mM NaCl, and 1% Triton-X. This buffer is referred to as sample buffer throughout the whole study. For RNA samples, a modified RNA sample buffer was used, consisting of sterile filtered sample buffer, modified with 0.1% RNase inhibitor, 1 mM DTT, and 1% carrier RNA.

**Table S1**. DNA sequences of the MAC circuit.

| Strand (corresponding domains) | Sequence |
|---|---|
| Target/Amplifier 2 [NC_045512.2, 21029-21072] (3+2+1) | 5'-GGG ATC TCA TTA TTA GTG ATA TGT ACG ACC CTA AGA CTA AAA A-3' |
| 27nt Target [NC_045512.2, 21029-21072] (3+2+1) | 5'-GGG ATC TCA TTA TTA GTG ATA TGT ACG-3' |
| 37nt Target [NC_045512.2, 21029-21072] (3+2+1) | 5'-GGG ATC TCA TTA TTA GTG ATA TGT ACG ACC CTA AGA C -3' |
| 59nt Target | 5'-TAA TAA AT GGG ATC TCA TTA TTA GTG ATA TGT ACG ACC CTA AGA CTA AAA A TGT TAC AA-3' |

| | |
|---|---|
| [NC_045512.2, 21029-21072] (3+2+1) | |
| Label DNA 1 (1) | 5‘-A CC CTA AGA CTA AAA A (AAAAA AAAAA)-3‘ + Biotin-TEG |
| Label DNA 2 (4) | Biotin-TEG + 5‘-(AAAAA AAAAA) CCC ACC CAC TTC CAC CCA-3‘ |
| Label DNA 3 (5) | 5‘-CC TTT TTC AAT ACC CTA CC (AAAAA AAAAA)-3‘ + Biotin-TEG |
| Label DNA 4 (7) | Biotin-TEG + 5‘-(AAAAA AAAAA) TAA TTG TAT GTG TGT AGC-3‘ |
| Amplifier 1 (2+6+5) | 5‘-CAT TAT TAG TGA TAT GTA CG TTT CTC CCC TTT TTC AAT ACC CTA CC-3’ |
| Substrate 1 (1*+2*+3*+4*) | 5‘-TTT TTA GTC TTA GGG TCG TAC ATA TCA CTA ATA ATG AGA TCC CTGG GTG GAA GTG GGT GGG-3‘ |
| Substrate 2 (5*+6*+2*+7*) | 5‘-GGT AGG GTA TTG AAA AAG GGG AGA AA CGT ACA TAT CAC TAA TAA TGG CTA CAC ACA TAC AAT TA-3‘ |
| 1 mismatch | 5‘-GGC ATC TCA TTA TTA GTG ATA TGT ACG ACC CTA AGA CTA AAA A-3‘ |
| 2 mismatches | 5‘-GGC ATC TCA TTA TTA CTG ATA TGT ACG ACC CTA AGA CTA AAA A-3‘ |
| 6 mismatches | 5‘-GCC TTC TCA TTA TTA GTG ATA TGT TGC ACC CTA AGA CTA AAA A-3‘ |

**Table S2**. RNA sequences of the MAC circuit.

| Strand (corresponding domains) | Sequence |
|---|---|
| Target [NC_045512.2, 21029-21072] (3+2+1) | 5’-GGG AUC UCA UUA UUA GUG AUA UGU ACG ACC CUA AGA CUA AAA A-3’ |
| 27nt Target [NC_045512.2, 21029-21072] (3+2+1) | 5’-GGG AUC UCA UUA UUA GUG AUA UGU ACG-3’ |
| 37nt Target [NC_045512.2, 21029-21072] (3+2+1) | 5’-GGG AUC UCA UUA UUA GUG AUA UGU ACG ACC CUA AGA C -3’ |
| 59nt Target [NC_045512.2, 21029-21072] (3+2+1) | 5’-UAA UAA AU GGG AUC UCA UUA UUA GUG AUA UGU ACG ACC CUA AGA CUA AAA A UGU UAC AA-3’ |

**Sample preparation protocols:**

**Preparation of adapted cDNA substrates**

To prepare the adapted cDNA substrate, cDNA (Substrate 1 or Substrate 2, 20 µl, 20 µM) and the corresponding amplifier ssDNA (Amplifier 2 or Amplifier 1, 20 µl, 40 µM) were mixed in 10 mM TRIS-HCl buffer solutions (pH 7.7, 150 mM NaCl, 0.01% Tween 20). The mixtures were annealed at 95°C for 5 min and cooled to room temperature within 120 min using a thermocycler. In the following, adapted substrate 1 refers to Substrate 1 + Amplifier 2 and adapted substrate 2 refers to Substrate 2 + Amplifier 1.

**DNA labelling of BNF80**

BNF80 particles were functionalized with Label DNA via streptavidin-biotin binding reaction. In total, four base solutions were prepared for each Label DNA. In the following, base solution 1 will refer to BNF80 + Label DNA 1, base solution 2 will refer to BNF80 + Label DNA 2, base solution 3 will refer to BNF80 + Label DNA 3 and base solution 4 will refer to BNF80 + Label DNA 4.

The base solutions were prepared with a total volume of 18.75 µl. Prior to functionalization, the BNF80 stock solution was sonicated for 30 min to dissolve possible aggregates that could result from storage. Then, a 0.375 µl of BNF80 stock solution ($c_{Fe}$= 5.5 g/l, 20 nM particles, streptavidin-coated) were mixed with 17.9 µl sample buffer and incubated with 0.46 µl (1 pmol/µl) of Label DNA. The incubation was done in a thermomixer at 25°C and 10 min cycles (5 min mixing at 650 rpm and 5 min at 0 rpm) over a course of 24 h. After functionalization, the DNA-labelled BNF80 particles were purified from excess biotinylated Label DNA by centrifugation at 3000 rcf and 21°C for 30 min. The supernatant was removed and fresh sample buffer was added. The washing step has been repeated two times. Afterwards, the volume of the base solutions has been adjusted to its initial volume of 18.75 µl. The base solutions are then used for the preparation of responsive magnetic clusters (RMCs).

**Preparation of RMCs**

To form responsive magnetic clusters (RMCs), the base solutions consisting of DNA-labelled BNF80-MNPs are tethered into clusters via adapted substrate. Final RMC solutions consist of 0.4 nM BNF80 and 3 nM of each adapted substrate in a 75 µl TRIS-HCl buffer.

To form RMC 1, 18.75 µl of base solutions 1 and 2 were mixed with 2.25 µl (0.1 pmol/µl) adapted substrate 1 and the volume was adjusted to 75 µl with sample buffer. To form RMC 2, 18.75 µl of base solutions 3 and 4 were mixed with 2.25 µl (0.1 pmol/µl) adapted substrate 2 and the volume was adjusted to 75 µl with sample buffer. For the hybridization process, the samples were incubated in a thermomixer at 25°C and 10 min cycles (5 min mixing at 650 rpm and 5 min at 0 rpm) over a course of 24 h. After incubation, the RMCs were purified by centrifugation two times at 1000 rcf for 5 min at 4°C. Afterwards, the volume has been concentrated six times for further processing.

We noted that cluster formation and washing depend on the total incubation volume. For the sake of reproducibility, the RMCs were always prepared for five MAC assays. Consequently, the total incubation volume for RMCs added up to 375 µl. After incubation, the volume has been reduced to 62.5 µl, sufficient for five MAC assays.

**MAC assays for DNA detection**

In a typical DNA assay, the desired amount of target DNA was first diluted to 7.5 µl with the sample buffer. Then, 12.5 µl of each RMC system was added to the target DNA solution, meaning a total incubation volume of 32.5 µl. The sample was incubated at 25°C for 24 h using thermomixer in 10 min (each cycle: 2.5 min shaking at 1400 rpm and 7.5 min at 0 rpm). The volume was adjusted to 75 µl before assay read-out.

**MAC assays for RNA detection**

For RNA, the assay sample buffer was sterile filtered and then modified with 0.1% RNase inhibitor, 1 mM DTT and 1% carrier RNA. After purification of RMCs, the volume was adjusted using the RNA sample buffer.

For an RNA assay, the RNA target was diluted in 7.5 µl RNA sample buffer. Then, 12.5 µl of each RMC system was added to the target for a total incubation volume of 32.5 µl. The sample was incubated at 25°C for 24 h using a thermomixer in 10 min cycle with 2.5 min of shaking at 1400 rpm and 7.5 min at 0 rpm. The volume was adjusted to 75 µl before measuring.

**Non-amplifying magnetic circuit**

The non-amplifying assays were performed in the same way as the MAC assays with the only difference that here only RMC 1 system (25 µl) was added to the target solution. The sample volume and incubation conditions were kept the same as the MAC assays.

**Characterization techniques:**

**Magnetic Particle Spectroscopy (MPS)**

Custom-built MPS setup was used, operating with an AC magnetic field amplitude of 15 mT and an excitation frequency of 590 Hz. The MAC assays were performed on 60 µl of the assay mixture (refer sections on assay preparation). A blank sample (60 µl sample buffer) was measured before each assay sample. To evaluate the assay outcome, the harmonics ratio HR53 between $5^{th}$ and $3^{rd}$ harmonic is calculated, eliminating the influence of particle concentration on the assay result [1][2].

**Magnetic Property Measurement System (MPMS)**

Magnetic Property Measurement System (MPMS-3) from Quantum Design was used to measure the magnetization hysteresis loops of BNF-MNPs in different states. The magnetization measurements were performed in the vibrating-sample magnetometry (VSM) mode at magnetic fields between +7 and -7 Tesla. In a typical measurement, a 15 µl of BNF80 at a particle concentration of 0.2 nM was freeze-dried in 15 µl of mannitol in designated capsule.

**Dynamic Light Scattering (DLS)**

DLS was utilized to measure the hydrodynamic size of particles. In a typical measurement, a 50 µl of particles (BNF-MNPs, RMCs, and post-MAC) was diluted ten times in 500 µl of sample buffer and measured by a Nano ZetaSizer from Malvern Panalytical in 173° backscattered measurement mode. Malvern ZetaSizer operates at a wavelength of 632 nm.

**Alternating current susceptibility (ACS)**

A fluxgate sensor-based system was used to measure the ACS spectra of the BNF80 particles. The sample is exposed to an ac magnetic field with an amplitude of 0.5 mT and varying excitation frequencies, ranging from 1 Hz to 3 kHz [3]. The measurements were performed at room temperature on a 150 µl of particle suspension with a typical particle concentration of 0.2 nM.

**Scanning Transmission Electron Microscopy (STEM)**

STEM measurements were performed using a Helios™ 5 UX Dual Beam Focus Ion Beam-Scanning Electron Microscope (Thermo Fisher Scientific) in bright field transmission mode. To prepare a sample, 10 ul of a diluted sample was pipetted on a TEM grid (formvar-carbon coated copper grids with the mesh size of 300). After 5 min of incubation, the remaining suspension was removed by using filter paper. The grid was then left to thoroughly dry under fume hood.

**Fluxgate magnetorelaxometry (MRX)**

MRX measurements were performed using a custom-built setup. MNPs were first exposed to a magnetic field of 2 mT for 2 s. Right after switching the field off, the demagnetization process of the sample due to Brown and Néel relaxation of the particles is recorded for 1.5 s [4].

**Characterization of magnetic nanoparticles:**

**Static properties of BNF80**

STEM micrographs of BNF80-MNPsare shown in figure S1a and S2b. Figure S1a shows multiple BNF80 particles evenly distributed on the grid, which vary in size. Figure S1b depicts two typical particles of different shapes, that are neither of cubic nor spherical form. The BNF80 particles consist of multiple magnetite crystallites (dark grey) that are encapsulated within a hydroxyethyl starch shell (light grey), which holds the crystallites in place and provides a conjugation chemistry for streptavidin functionalization. The size distribution of the BNF80 particles and their crystallites (Figure S1c) were determined from the analysis of STEM images using the image analysis program ImageJ (https://imagej.nih.gov/ij/). The diameter of the single crystallites was estimated to be $d_{c,STEM}$ = 15.1 nm ± 3.5 nm, while the average diameter of the BNF80 particles was calculated to be $d_{h,STEM}$ = 116.7 nm ± 47.1 nm. The particle hydrodynamic size was measured by dynamic light scattering (DLS) method (Figure S1d). The average BNF80 hydrodynamic size was calculated to be $d_{DLS}$ = 138.9 nm ± 50.3 nm. BNF80 particles possess a broad hydrodynamic size distribution. Thorough vortexing of the BNF80 base solution is crucial to achieve reproducible samples.

The BNF80 particles were further analyzed via an magnetic property measurement system (MPMS-3, Quantum Design). The measured data (not shown) was analyzed using a fit routine developed by [5]. The saturation magnetization of the BNF80 particles was determined to be $M_{S,mass}$ = 89.89 $Am^2/kg_{Fe}$, which can be converted to $M_{S,vol}$ = 362.2 kA/m with the density of magnetite ρ = 5.18 $g/cm^3$ and the ratio of magnetite to the MNP of 1.2857 $g_{Magnetite}/g_{MNP}$ [6].

**Dynamic properties of BNF80**

MRX measurements were carried out on both liquid and freeze-dried particles. The results are shown in Figure S1d. The magnetization drops significantly faster for the BNF80 particles in suspension compared to the immobilized BNF80 particles, demonstrating that the Brownian relaxation time is significantly faster than the Néel relaxation time at room temperature. Due to the Brownian relaxation dominating the effective relaxation time, the BNF80 particles are suitable for immunoassay applications, which are based on the change of the hydrodynamic volume of the particle system upon binding.

Figure S1f depicts the ACS spectrum of the BNF80 particles normalized the initial static susceptibility $\chi_0$. The particle size was determined to be $d_{h,ACS}$ = 116.7 nm ± 29.1 nm by fitting the imaginary part χ'' peak of the ACS spectrum to the Debye model using a fit routine written in MATLAB [4]. This corresponds to the Brownian relaxation time constant of $\tau_{B,ACS}$ = 612.8 µs. The following parameters were set as fix parameters: particle effective core size of $d_{c,eff}$ = 39.7 nm, viscosity of the suspending medium η = $10^{-3}$ Pa·s, temperature T = 295 K, and $M_S$ = 362.2 kA/m. The average Brownian relaxation time $\tau_B$ can easily be estimated by plugging the frequency for which $\chi''/\chi_0$ peaks $f_{Peak}$ = 260 Hz into $2\pi \times f_{peak} \cdot \tau_B = 1$. This results in $\tau_B$ = 612.4 µs, matching the value extracted from the fit.

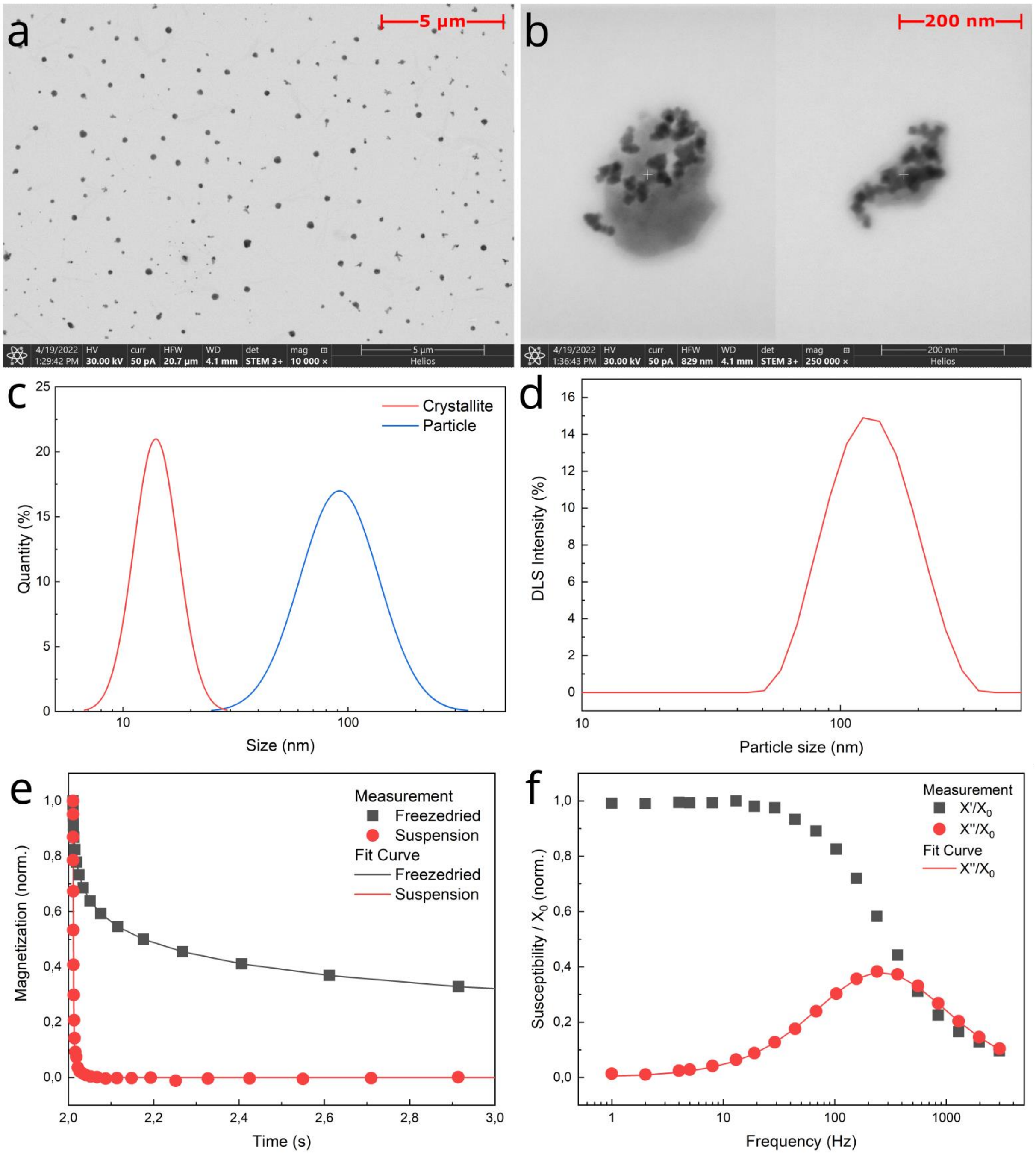


**Figure S1**. Measurement results for the characterization of BNF80-MNPs, a) STEM image of evenly distributed MNPs on a copper-coated TEM grid. b) STEM images of two different BNFs. c) Size distributions (crystallites and particle size) based on the STEM images, determined using the ImageJ image analysis software. d) Intensity weighted size distribution measured with DLS. e) MRX results of BNF80 both in suspension and freeze-dried in mannitol matrix, f) ACS results normalized to the initial static susceptibility $\chi_0$.

**Normalized MAC assay results**

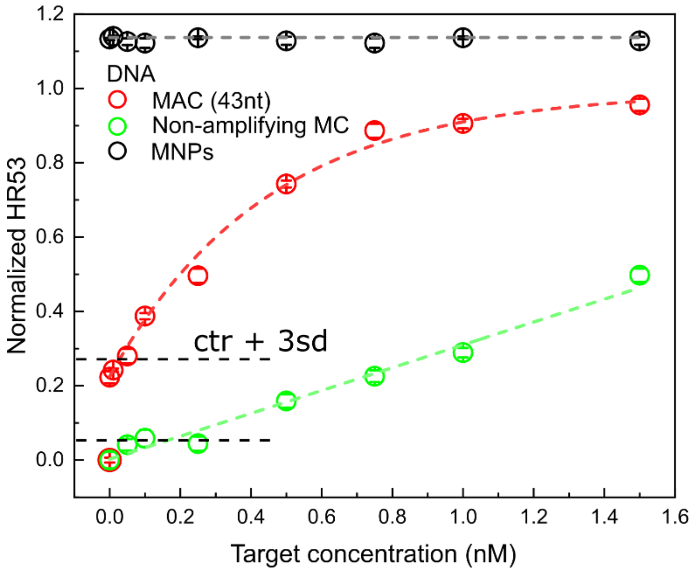


**Figure S2**. Normalized harmonics ratio as a function of target concentration. The harmonics ratio was normalized as follows: $(HR53_c - HR53_{min})/(HR53_{max} - HR53_{min})$, where $HR53_c$ is the harmonics ratio at a target concentration c and $HR53_{min}$ and $HR53_{max}$ are the harmonics ratios at 0 nM and 10 nM target concentration, respectively.

**Non-amplifying MC assay**

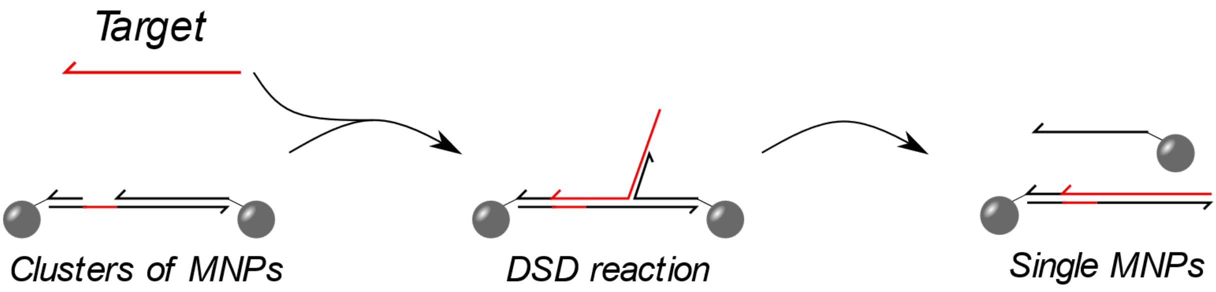


**Figure S3.** Non-amplifying assay concept based on the DSD reaction. The cDNA possesses a toehold, to which the target sequence can bind and subsequently displace the label DNA. Single MNPs as output are generated.

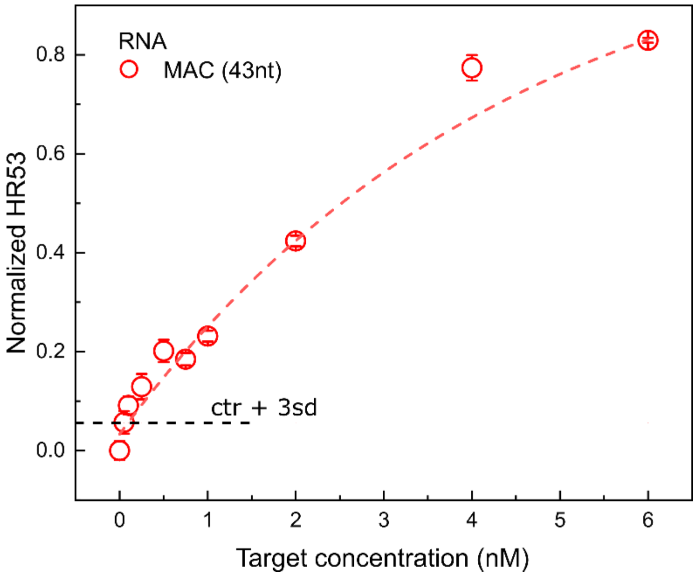


**Figure S4** Normalized harmonics ratio as a function of target concentration for the 43 nt RNA target (main text Fig. 3). The harmonics ratio was normalized as follows: $(HR53_c\text{-}HR53_{min})/(HR53_{max}\text{-}HR53_{min})$, where $HR53_c$ is the harmonics ratio at a target concentration c and $HR53_{min}$ and $HR53_{max}$ are the harmonics ratios at 0 nM and 10 nM target concentration, respectively.

**REFERENCES:**


[1] S. Draack *et al.*, “Multiparametric Magnetic Particle Spectroscopy of CoFe2O4 Nanoparticles in Viscous Media,” *Journal of Physical Chemistry C*, vol. 123, no. 11, pp. 6787–6801, Mar. 2019, doi: 10.1021/acs.jpcc.8b10763.

[2] A. M. Rauwerdink and J. B. Weaver, “Viscous effects on nanoparticle magnetization harmonics,” *J Magn Magn Mater*, vol. 322, no. 6, pp. 609–613, Mar. 2010, doi: 10.1016/j.jmmm.2009.10.024.

[3] J. Dieckhoff, A. Lak, M. Schilling, and F. Ludwig, “Protein detection with magnetic nanoparticles in a rotating magnetic field,” *J Appl Phys*, vol. 115, no. 2, Jan. 2014, doi: 10.1063/1.4861032.

[4] E. Heim, F. Ludwig, and M. Schilling, “Binding assays with streptavidin-functionalized superparamagnetic nanoparticles and biotinylated analytes using fluxgate magnetorelaxometry,” *J Magn Magn Mater*, vol. 321, no. 10, pp. 1628–1631, May 2009, doi: 10.1016/j.jmmm.2009.02.101.

[5] P. Bender *et al.*, “Relating Magnetic Properties and High Hyperthermia Performance of Iron Oxide Nanoflowers,” *The Journal of Physical Chemistry C*, vol. 122, no. 5, pp. 3068–3077, Feb. 2018, doi: 10.1021/acs.jpcc.7b11255.

[6] F. Ahrentorp *et al.*, “Effective particle magnetic moment of multi-core particles,” *J Magn Magn Mater*, vol. 380, pp. 221–226, Apr. 2015, doi: 10.1016/j.jmmm.2014.09.070.
